# Masking effects on *Iso*-valeric Acid Recognition by Sub-threshold Odor Mixture

**DOI:** 10.1101/2022.10.13.512096

**Authors:** J. Huang, J. Lin, R. Yueng, S. Wu, L. Solla, T. Acree

## Abstract

Masking unpleasant odors with high levels of pleasant-smelling odorants is an ancient practice that has evolved into many enterprises, from perfumery to consumer products. However, effective odor masking turns out to be idiosyncratic and impermanent. Here, we used Sniff Olfactometry (SO)(Rochelle et al., 2017; Wyckoff & Acree, 2017) to investigate the psychophysics of masking during 70ms-stimulations with mixtures of the mal-odorant *iso*-valeric Acid (IVA) and different masking agents. IVA is a component of human sweat that can dominate its smell, and is often described in unpleasant terms, e.g., “gym locker”, “smelly feet”, “dirty clothes”, etc. Conventionally, high concentrations of positive smelling odorants are used to reduce the unpleasantness of IVA in clothing or environments contaminated with IVA. To investigate the masking effects of sub-threshold levels of masking agents (neohivernal, geraniol, florhydral, decanal, *iso*-longifolanone, methyl *iso*-eugenol, and *s-*limonene) on IVA, we used SO to measure the probability of recognizing IVA after 70ms stimulations with headspaces containing mixtures of super-threshold concentrations of IVA and sub-threshold concentrations of IVA-suppressors for 9 subjects. On average, the single masking agent could decrease IVA-recognition probability by 14% to 72%, and a subthreshold odor mixture consisting of 6 masking agents decreased IVA recognition by 96%.

## Introduction

### Background

When consumers preferentially choose scented products over their unscented versions, it is partly to mitigate malodor often associated with the consumer products (Herz et al., 2022). Using a pleasant odor to cover malodor has been shown to reduce the negative impact on well-being that malodors can produce (Dalton et al., 2020). Traditionally, strong pleasant odors are used to cover the unpleasant odors: eg. using suprathreshold level of citronellal, limonene and citral to cover dimethyl sulfide citronelle (Osada et al., 2013). However, a series of health hazards including sensory irritation, respiratory symptoms, and dysfunction of lungs have been associated with exposure to high levels of fragrance (Kim S, 2015; Steinemann, 2016). To limit the amounts of odorants exposed to consumers, we studied the use of peri- and/or sub-threshold amounts of masking agents could mask IVA while the masking agents remained undetectable or barely detectable. Although definitions of odor masking (i.e., the modification of perceived odor quality to make it more acceptable) and odor counteraction (i.e., the reduction of perceived intensity) are quite established. (Bell, 1987; Jones, 1964; Laing & Francis, 1989; Laing, 1984; Oka, Omura, et al., 2004; Osada et al., 2013) Little work has been done to address the concentration of masking agents used to mask the malodor. Herein, we define “**Odor Covering**” as using supra-threshold amount of strong pleasant odorants to cover the malodor (counteraction and masking), “**Odor Masking**” as using barely detectable amount (sub-threshold or peri-threshold) pleasant odorants to make malodor unrecognizable or slightly pleasant.

### Neurological and Pharmacological Rationale

Odor perception has been described as a process of encoding and decoding in which odor encoding starts with the binding of odorants to specific sets of olfactory receptors (ORs) in the olfactory epithelium (Buck, 1996; Kajiya et al., 2001; Malnic B, 1999). These odorants either stimulate (Agonists) or inhibit some ORs (Antagonists) (Marc Spehr, 2003; Oka, Nakamura, et al., 2004; Oka, Omura, et al., 2004; Ricardo C. Araneda, 2000) to form a unique ORs coding which is transduced to the olfactory bulbs (OBs) in glomeruli (Buck, 2000), where the signals are further transduced and decoded in the central nervous system (Strutz et al., 2014). It well established that human odor perceptions are related to specific receptor activations (Keller et al., 2007; Menashe et al., 2007) and inhibition (Pfister et al., 2020; Reddy et al., 2018). Some preliminary research (Aya Kato, 2015; KAO et al., 2019) have shown odorants that are antagonistic to malodors can reduce the intensity of malodor when mixed. In this study, total of 7 perfume raw materials (PRMs) were selected as the potential masking agents against IVA: **1**. Neohivernal (Neo), a PRM reported to reduce IVA intensity (Stacy Renee Hertenstein et al., 2017); **2**. Florhydral (Flo), a PRM reported to be an antagonist to the IVA receptor OR51E1 (Aya Kato, 2015); **3**. Methyl *Iso*-eugenol (Met), Decanal (Dec), *Iso*-longifolanone (Long), Geraniol (Ger) and *s*-Limonene, PRMs traditionally used to reduce malodor but did not interact with OR51E1 (Bushdid et al., 2018; Halperin Kuhns et al., 2019; Saito et al., 2009; Sean M. Wetterer, 2015).

### Experiment Design Rationale

Since odor coding is concentration dependent for both single odorant and mixture (Kajiya et al., 2001; Xu et al., 2020), and humans respond to sub-threshold odorants (Hummel et al., 2013), we hypothesize that sub-threshold IVA odor-suppressors can, to some extent, mask human IVA perception. To investigate the effects of sub-threshold IVA odor-suppressors, we used SO to compare IVA detection probability after 70ms stimulations with headspaces containing mixtures of super-threshold concentrations of IVA with and without sub-threshold concentrations of IVA-suppressors. Although previous research failed to show interactions between low intensity odorants that promoted or inhibit each other (Laing, 1984), we have explored whether a mixture of sub-threshold concentrations of masking agents can significantly mask IVA while remaining undetectable (Rochelle et al., 2017).

### Materials

#### Chemicals

Polyethylene Glycol 400 (PEG400): CAS Registry No. 9002-88-4, JT Baker^®^, Avantor Performance Materials, Inc, (>99.5%).

90% Deionized Water: carbon filtered deionized water.

Charcoal powder: CAS Registry No. 7440-44-0, Activated Charcoal Norit^®^ Norit^®^ SX2, powder, from peat, multi-purpose activated charcoal, steam activated, and acid washed.

*Iso*-valeric Acid (IVA): CAS Registry No. 503-74-2, Sigma Aldrich (> 99%). Geraniol (Ger): CAS Registry No. 106-24-1, Vigon International, Inc (> 100%). Neohivernal (Neo): CAS Registry No. 300371-33-9, Firmenich Inc, (> 99%).

Florhydral (Flo): CAS Registry No.125109-85-5, Givaudan Fragrances Corp. (> 90%). Decanal (Dec): CAS Registry No. 112-31-2, Givaudan Fragrances Corp. (> 90%).

*Iso*longifolanone (Long): CAS Registry No. 14727-47-0, Vigon International, Inc (100%). Methyl *Iso*-eugenol (Met): CAS Registry No. 6379-72-2, Givaudan Fragrances Corp. (> 90%). *S*-Limonene (Lim): CAS Registry No. 5989-54-8, Sigma Aldrich (> 99%).

### Subjects

9 subjects including 7 females and 2 males, all students from Cornell University (22-27 years old), were tested. Subjects participated were screened to make sure they 1) do not have a stuffy nose before each session; 2) do not have post-COVID anosmia; 3) could smell all the compounds.

### Software

The experiments were automated using PsychoPy® (v2021.2.3) (Peirce et al., 2019). Data analysis was executed using R (version 4.1.3 – “One Push-Up”) (R-Core-Team, 2022). See supplemental.

## Methods

### Sample Preparation

#### Stock Solution Preparation

1000 PPM stock solution for each odorant in PEG 400 was prepared. 0.1g of each odorant was added to 100mL amber bottle followed by 100mL PEG 400 addition.

#### 10% PEG – Water Solution Deodorization

400mL of PEG 400 was added 3600mL of DI water. The mixture solution was added 20g of Charcoal powder and vigorously mixed. The mixture was set for 2 days and performed vacuum filtration to remove the charcoal powder to obtain the deodorized 10% PEG – water solution.

#### Test Sample Solution Preparation

Each odorant was diluted to different concentrations to make 50mL 10% PEG water solution. The solutions were prepared 1 day before the experiment, mixed on a shaker overnight, and transferred to a 250mL Teflon bottle 10mins before experiments.

### Threshold Determination

#### Descriptor Determination

Each odorant was assigned a consensus descriptor that matched its odor characteristics as follows: IVA (Stinky Feet), neohivernal (Clean Laundry), geraniol (Rosey), florhydral (Floral), *iso-*longifolanone (Woody), decanal (Soapy), methyl *iso*-eugenol (Clove) and *s*-limonene (Citrus).

#### Concentrations Within a Triad

Three concentrations were determined by the following rules:

1. Mutual difference between concentrations were greater than (ΔC/C ≧0.33)
2. Two test runs were performed on lab members to be assured that the three concentrations generate a robust logistic function for odor threshold.

#### Conditioning and training

Highest concentration we used in a test was used in the conditioning and training. Each subject sat down, adjusted the chair to the proper height and positioned their nose above the sniffing port before the PsychoPy® session began (Ni et al., 2022). Then, the subjects started the first trial and followed the instructions and cues shown on the until the trial ended in about 8 seconds. Prompts including “Click when you are ready”, “When you are ready to inhale, Click again”, “Inhale”, and “Exhale”. 700ms seconds after the cue to inhale is displayed the SO will puff a 15ml blast of headspace gas for 70ms. Then the monitor will prompt “This smell is XXX (descriptor)”. The subjects will repeat this process for up to 6 times to familiarize themselves with the odor. Subjects who were not able to smell this concentration were discontinued for the next session.

#### Pre-testing Session

The bottle containing solution used in the training session and the bottle containing blank sample (10% PEG and water) were used in this session. The procedure was the same as the training session until after the puff, a binary forced choice question “Did you smell xxx (descriptor for that odorant)?” was shown on the monitor and the subjects were required to choose “Yes”, or “No” to continue. The process was repeated for 5 puffs for each bottle; subjects were required to attain 90% accuracy on this trial before moving into the threshold measurement stage of the experiment.

#### Threshold Measurement

3 bottles containing ascending concentrations of an odorant, labeled 1 to 3 were puffed 4 times randomly each at each position in the triad (Supplemental). The probability of detecting the odor at designated concentration was plotted against log concentration and then fitted to a binary logistic model to yield a psychometric function. The recognition threshold was obtained by calculating the concentration where the detecting probability equals 0.5 (Ni et al., 2022; Wichmann, 2018).The threshold for IVA and 7 masking agents were measured for all 9 subjects. The threshold was measured for 1-3 times. Subjects were divided into 2 groups based on their IVA sensitivity. Subjects in Group 1 are sensitive to IVA (Recognition threshold < 4PPM) and subjects in Group 2 are moderately sensitive to IVA (Recognition threshold > 4PPM). Subjects who are hyposensitive to IVA were not used in the experiments.

### Masking Effect Determination

#### Masking IVA by subthreshold single odor mixture

To the three bottles of the SO triad were added: 50mL Supra-threshold amount of IVA (Bottle 1), 50mL mixture of Supra-threshold amount of IVA and sub-threshold level of first masking agent (Bottle 2), and 50mL mixture of Supra-threshold amount of IVA and sub-threshold level of another masking agent (Bottle 3). One experiment session includes 3 trials where each bottle was puffed 4 times (Supplemental Table 2) at each position on the triad (Supplemental Figure 1). After each puff, subjects were asked to answer “Yes” or “No” to the question: “Did you smell Stinky Feet (IVA)?” Each subject was asked to complete 2 sessions at each visit.

#### Masking IVA by sub-threshold amount of masking agent mixture

To three bottles on the SO triad were added: 50mL Supra-threshold amount of IVA (Bottle 1), 50mL mixture of Supra-threshold amount of IVA with sub-threshold level of masking agent mixture (Bottle 2), and 50mL blank sample contains 10% PEG in water solution (Bottle 3). For IVA-sensitive subjects, 10 PPM IVA was used, and for subjects having moderate IVA sensitivity, 15 PPM IVA was used. One experiment session includes 3 trials where each bottle was puffed 4 times randomly at each position on the triad (Figure 1). After each puff, subjects were asked to answer “Yes” or “No” to the question: “Did you smell Stinky Feet (IVA)?” Each subject was asked to complete 2 sessions at each visit.

**Figure 1.**
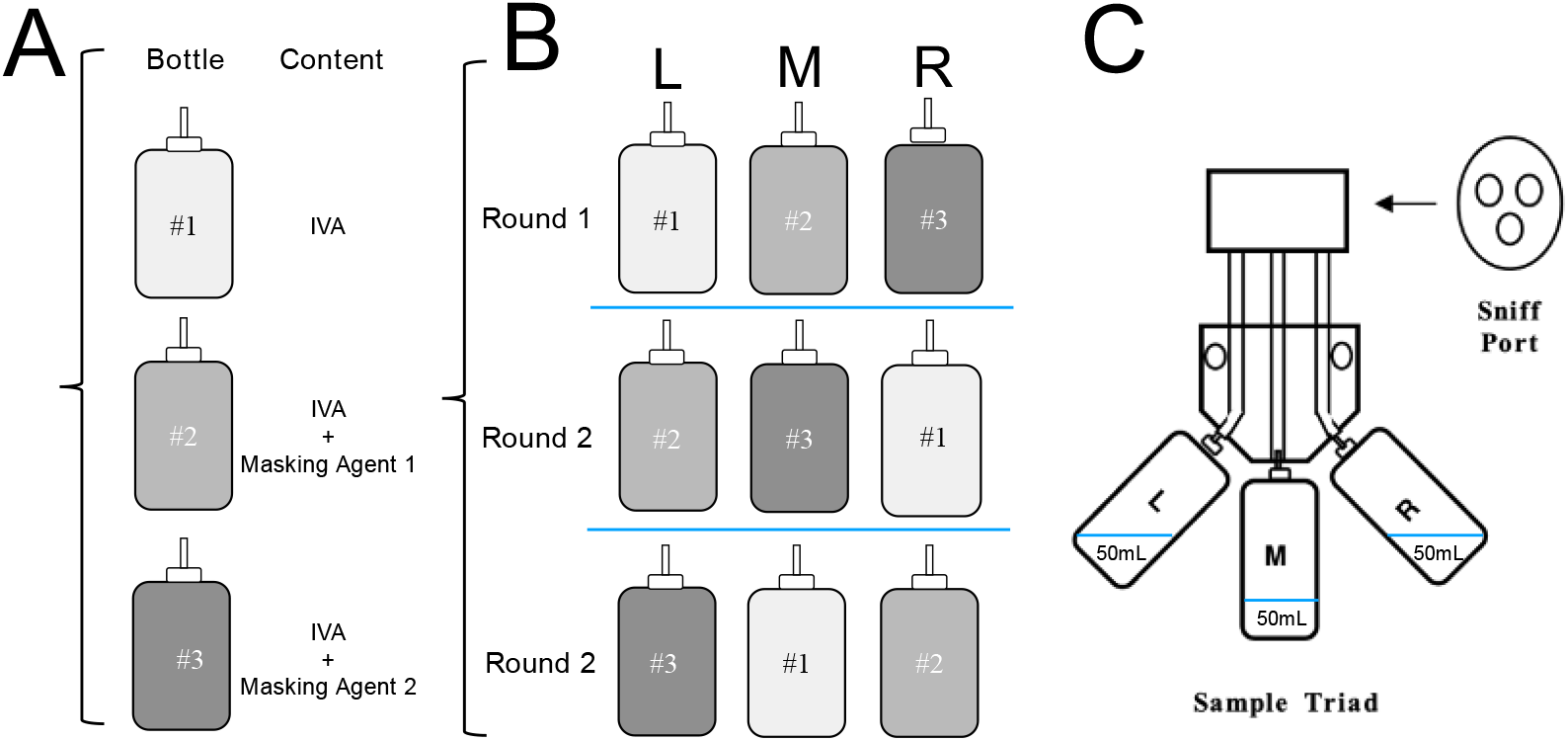
Figure1A shows the content of the solution in each bottle. Figure 1B shows 3 bottles was puffed 4 times randomly in one trial. For each experiment session, total of 3 trials were conducted, so that each bottle was puffed 4 times at each position on the triad shown in Figure 1B. Figure 1C displays the arrangement of bottles relative to the sniff port.

## Results

### Thresholds Measurement

Each subject’s thresholds for IVA and Masking Agents were measured using SO and the values were shown in supplement. Based on the IVA threshold we obtained, the subjects were divided into 2 subgroups: group 1 and group2. Group 1 includes subjects 1, 2, 3, 4, 5 who were sensitive to IVA (threshold < 4PPM), whereas Group 2 includes subjects 6, 7, 8, 9 who were moderately sensitive to IVA (threshold > 4PPM).

### Experiment Concentration Determination

For the subjects in group 1, 5PPM of IVA was used and for the subjects in group 2, 10PPM of IVA was used to make sure the IVA is at a detectable level but not overly strong. For each subject, concentrations at around half of masking agents’ detection thresholds were used (Figure 2A). Since the Lim1 concentrations for limonene were recognized by all the subjects, the experiment was repeated using 1 PPM limonene for everyone.

**Figure 2.**
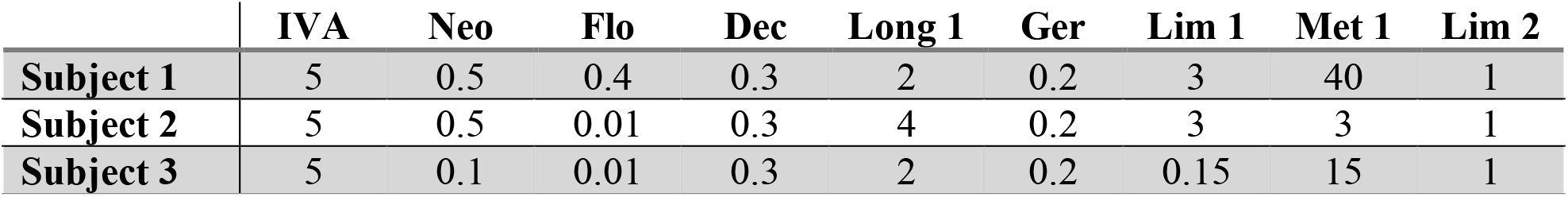
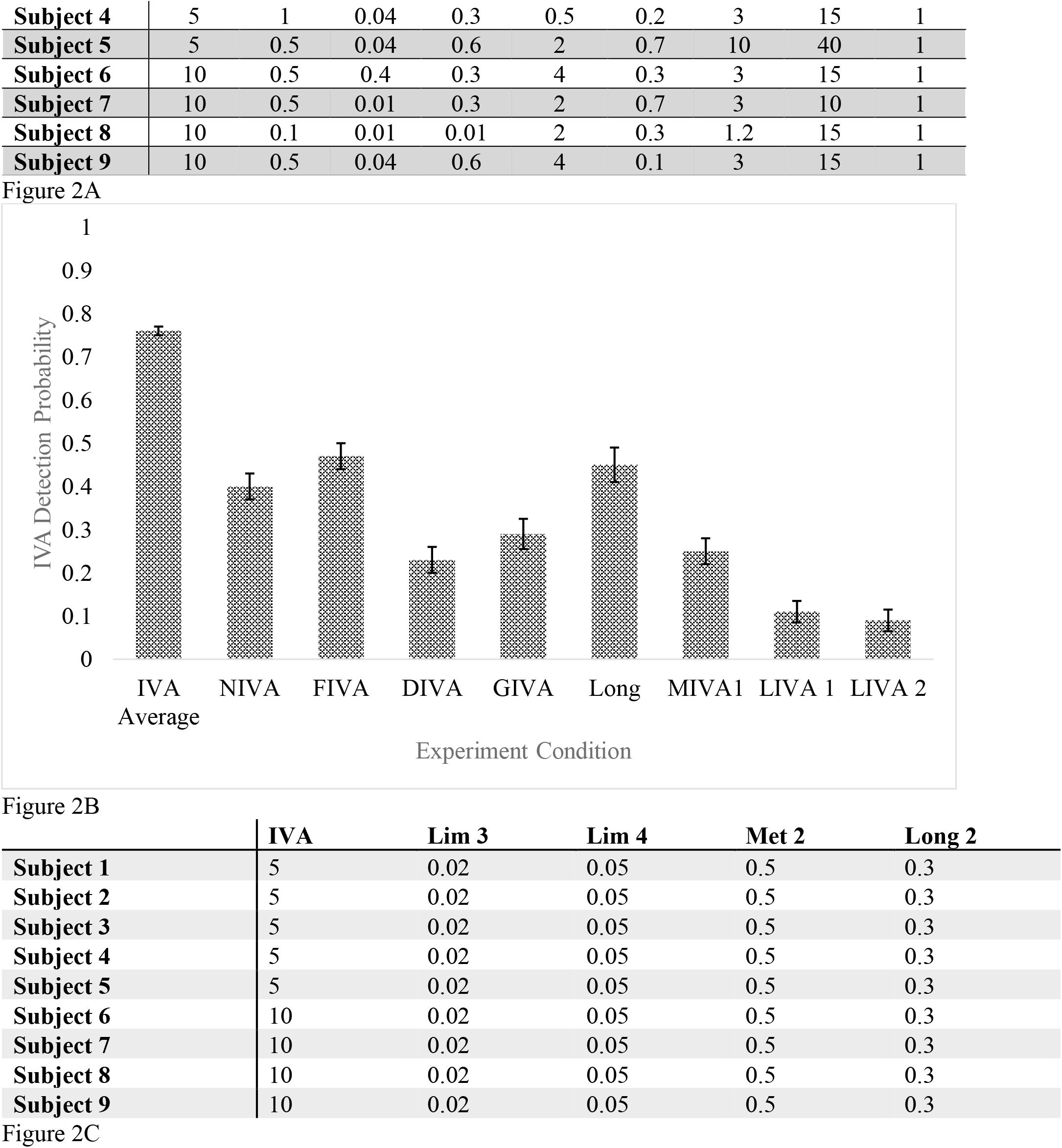
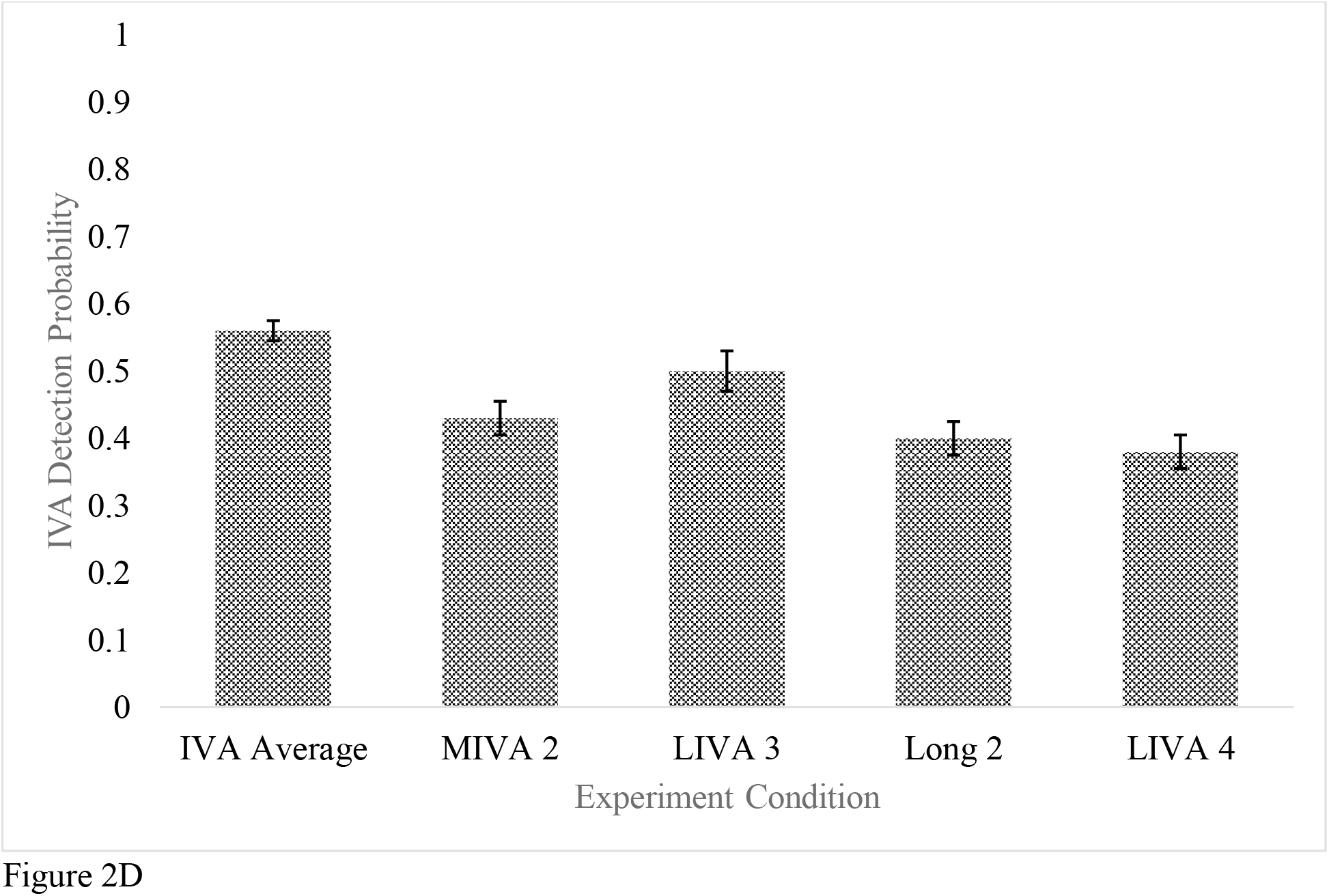
Figure 2A shows the amount of each PRM used in masking study in PPM. Figure 3B shows IVA Detection probability under all conditions. The IVA detection probabilities for the IVA were averaged to obtain the data for IVA average. NIVA stands for neohivernal + IVA, FIVA stands for florhydral + IVA, DIVA stands for decanal + IVA, GIVA stands for geraniol + IVA, Long stands for Iso-longifolanone + IVA. MIVA stands for methyl iso-eugenol + IVA. Lim stands for s-limonene + IVA. Figure 2C shows concentrations of each odorant used for the 2^nd^ round of IVA masking analysis. Figure 2D shows IVA detection probability with lowered masking agents’ concentrations for methyl iso-eugenol, Limonene and Isolongifolanone.

### Masking Effect Measurement 1

The IVA detection probability, P(IVA Detection) = Number of IVA stimuli detected / total number of IVA stimuli, was plotted against the experiment conditions (Figure 2B). After the SO experiment, subjects were asked if they were able to detect any other smell. Seven out of 9 were able to detect limonene at 1 PPM, thus 0.02 PPM and 0.05 PPM limonene were tested for their masking effects against IVA. Also, since the amount of methyl *iso-*eugenol in the mixture is more than the amount of IVA for some subjects, a lower concentration of methyl *iso-*eugenol (0.5PPM) was tested. (Figure 2C).

### Masking Effect Measurement 2

The IVA detection probability with lowered *s-*limonene, methyl *iso*-eugenol and *iso*-longifolanone concentrations were shown in Figure 2D. A Seemingly IVA habituation was observed at session 5 and 6, where the IVA detection probability for the pure IVA drops to around 50%.

### Masking Agents Mixture Determination

Every masking agent except for methyl *iso*-eugenol was used to make a 1.2 PPM masking agent containing 6 sub-threshold levels of masking agents. (Figure 3A) To solve the problem of IVA habituation, subjects in Group 1 were given 10PPM IVA while subjects in Group 2 were given 15PPM IVA. Thus the 3 triads contained a Blank (10% PEG), 10PPM or 15 PPM IVA, and 10PPM or 15PPM IVA plus 1.2ppm masking mixture.

**Figure 3.**
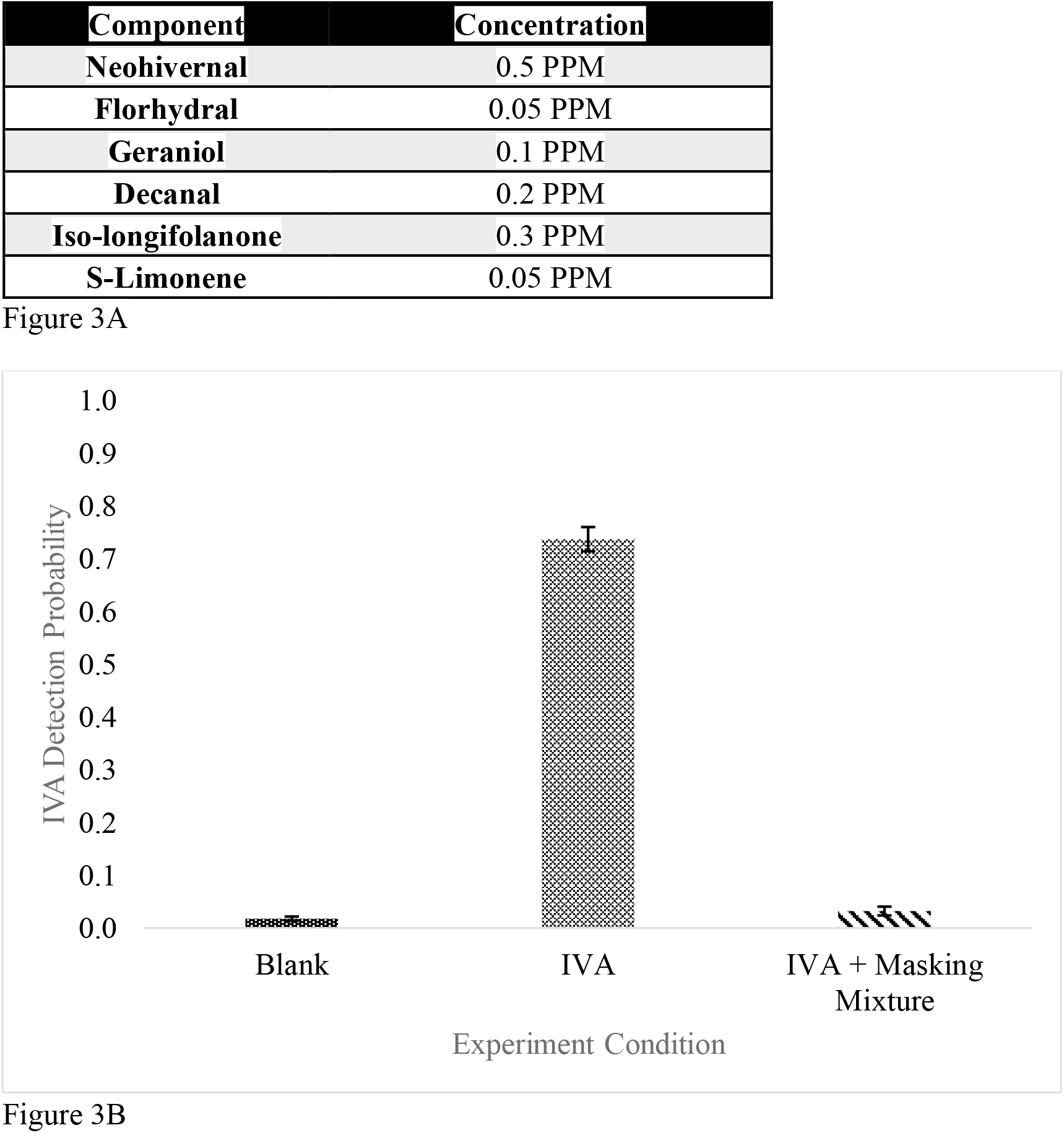
Figure 3A shows masking agent mixture components and their concentration. Figure 3B shows the plot of IVA detection probability for blank sample, 10PPM or 15PPM IVA, and 10PPM or 15PPM IVA with 1.2 PPM masking agent mixture, which clearly demonstrates the IVA was undetectable when mixing with the masking agent mixture.

### Masking Agent Mixture Masking Effects

As shown in Figure 3B, 8 out of 9 subjects were not able to perceive IVA when masked by a masking agent mixture, though not complete and the other subject showed a drastic decrease in IVA detection. 3 subjects reported no detectable smell for the mixture, while the other 3 subjects reported that when IVA was masked with the masking agent mixture, it possessed “a slight pleasant smell with touch of a clean and citrus note” and no specific odorants could be identified.

## Discussion

### Dose Dependence of Odor Masking

First, the masking capacity of each masking agent was calculated by Masking Capacity = (P_IVA_ – P_Masked IVA_)/P_IVA_. The masking capacities of 0.3PPM *iso*-longifolanone, 0.05PPM *s-*limonene and 0.5PPM methyl *iso-*eugenol along with the other masking agents were calculated. The masking capacity was plotted against the concentrations of masking agents used (Figure 4A). The masking concentration and masking capacity seem to demonstrate a dose-dependent effect: the higher concentration of masking agents used, the better masking capacity it possessed, except for methyl *iso-*eugenol for which high concentration does not convert into better masking capacity. Thus, a linear correlation plot between dose of masking agents (except methyl *iso-*eugenol) with their masking capacity is shown (Figure 4B). The Pearson Coefficient and p-value were calculated to be 0.51 and 0.85, indicating a weak linear trend between these two factors. The greater the dose of masking agents applied, the better the odor masking effect. However, in our case, we are considering both odor masking and odor covering within the same context, which could be the reason we observe a large p-value for the linear relationship. In the optimal cases, a better linear relationship could be established if we were just investigating odor masking or odor covering alone.

**Figure 4.**
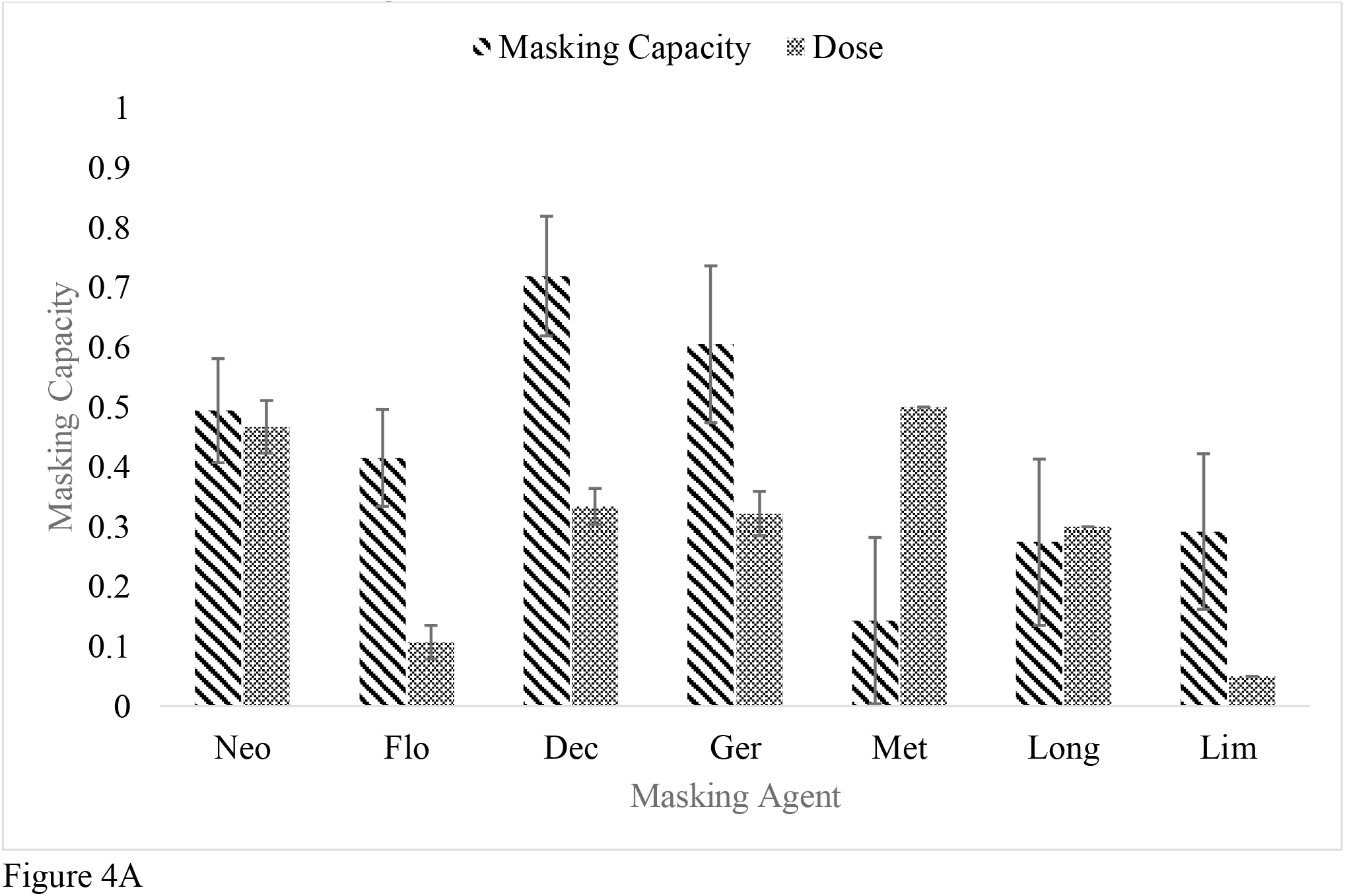
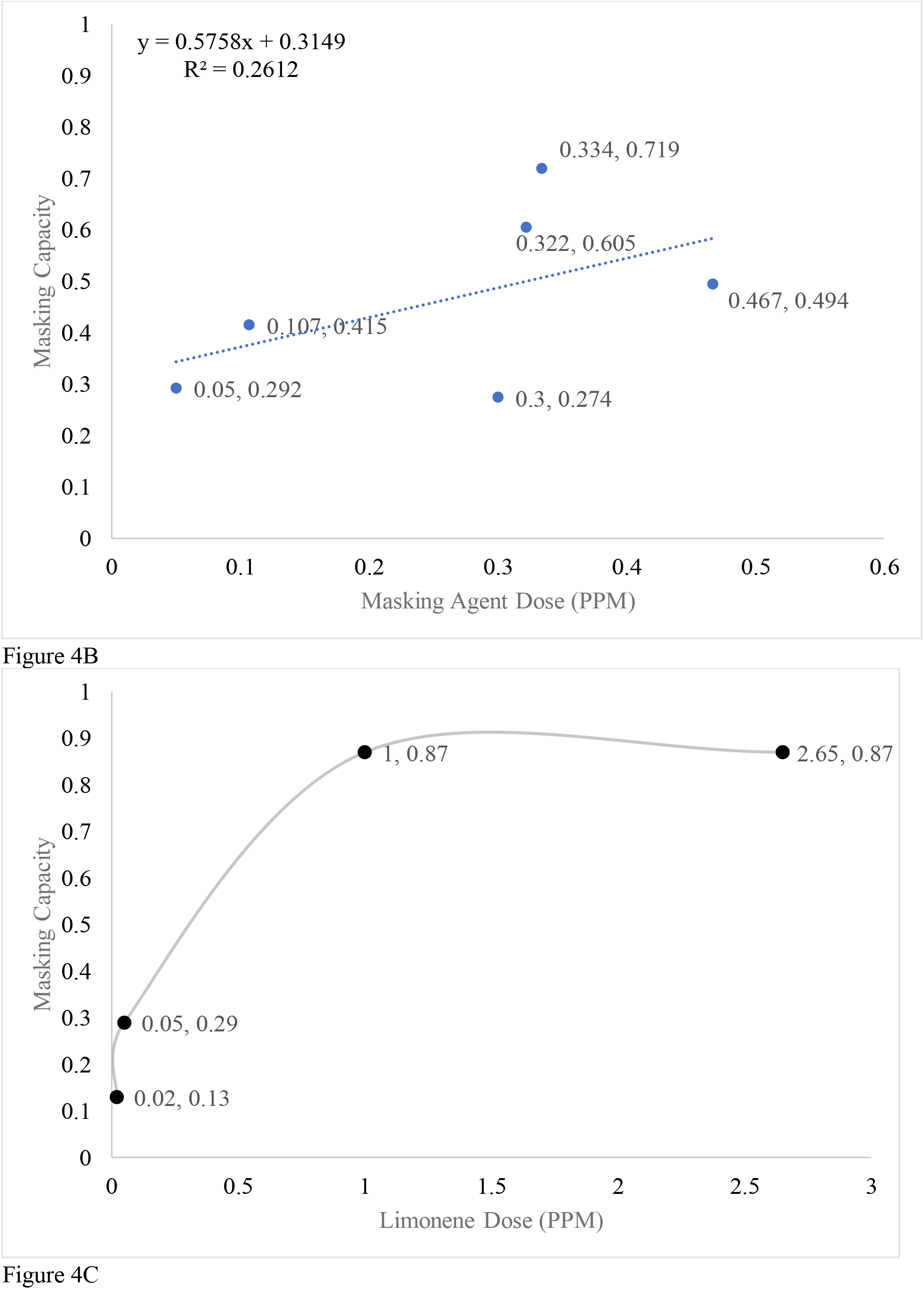
In figure 4A, IVA Masking Capacity was plotted against the dose of masking agents applied, showing the higher concentration of masking agents used, the better masking capacity it possessed. Figure 4B shows the linear correlation curve between dose of the masking agents and their corresponding masking capacities. 0.3PPM of Isolongifolanone and 0.05PPM of s-limonene were included while methyl iso-eugenol was excluded.). The Pearson Coefficient and p-value were calculated to be 0.51 and 0.85, which means there is a weak linear trend between these two factors. In Figure 4C, dose & masking capacity response curve was plotted for s-limonene at 0.02PPM, 0.05PPM, 1PPM and 2.65PPM, showing the dose and masking capacity follow a dose-dependence relationship until reaches a critical point where further increase in masking agents’ concentration does not convert to increase in masking capacity.

To further corroborate this phenomenon, *s-*limonene was used to study the dose-masking relationship. The overall masking capacity (both masking and covering) was plotted against the concentration of *s-*limonene used (Figure 4C). From 0.02PPM to 0.05 PPM, there was an increase in *s-*limonene masking capacity, while once it hits 1 PPM, the masking capacity of *s-* limonene seems to reach its maximum. Though for some subjects, 1PPM of *s-*limonene was able to completely mask the IVA, for others, the masking capacity did not show a significant increase using *s-*limonene above 1PPM. These results indicate that beyond a certain concentration further increases in dose does not increase to greater intensity. In summary, beyond a critical concentration ratio, masking does not increase as dose increases.

### Odor Masking and Odor Covering

In this experiment, we have defined “**Odor Covering**” as using supra-threshold amount of strong pleasant odorants to cover the malodor, “**Odor Masking**” as using barely detectable amount (sub-threshold or peri-threshold) pleasant odorants to make malodor unrecognizable or slightly pleasant. It has been reported that the total intensity of a mixture is less than the sum of the intensities of the components (Bell, 1987; Jones, 1964; Laing & Francis, 1989; Laing, 1984). Therefore when both odors are detectable, there is always mutual suppression (counteraction) effects that cause individual components in the mixture to become less intense (Cain & Drexler, 1974; Kittel et al., 2008; Kurtz et al., 2009; Kurtz et al., 2010; Weiss et al., 2012b). In case of Odor Covering, since the masking agents were detectable to subjects, both counteraction and odor masking effect were likely causing the decrease in IVA detection probability. To distinguish the difference between odor covering and odor masking, we compare the IVA detectability when subjects could and could not detect masking agents (Figure 5A). When subjects were able to detect the masking agents, the IVA detection probability dropped to 0.2; for subjects who were not able to detect the masking agents, the IVA detection probability stayed at 0.48. To further see the difference between odor covering and odor masking, the masking capacity of odor masking and odor covering were compared (Figure 5B). Figure 5B shows that odor masking is responsible for about 37% of decrease in IVA detection probability, while odor covering is responsible for 74% of decrease in IVA detection probability. The mutual suppression effect might be responsible for the difference in odor masking and odor covering, indicating that mutual suppression only happens when 2 odors are both detectable.

**Figure 5.**
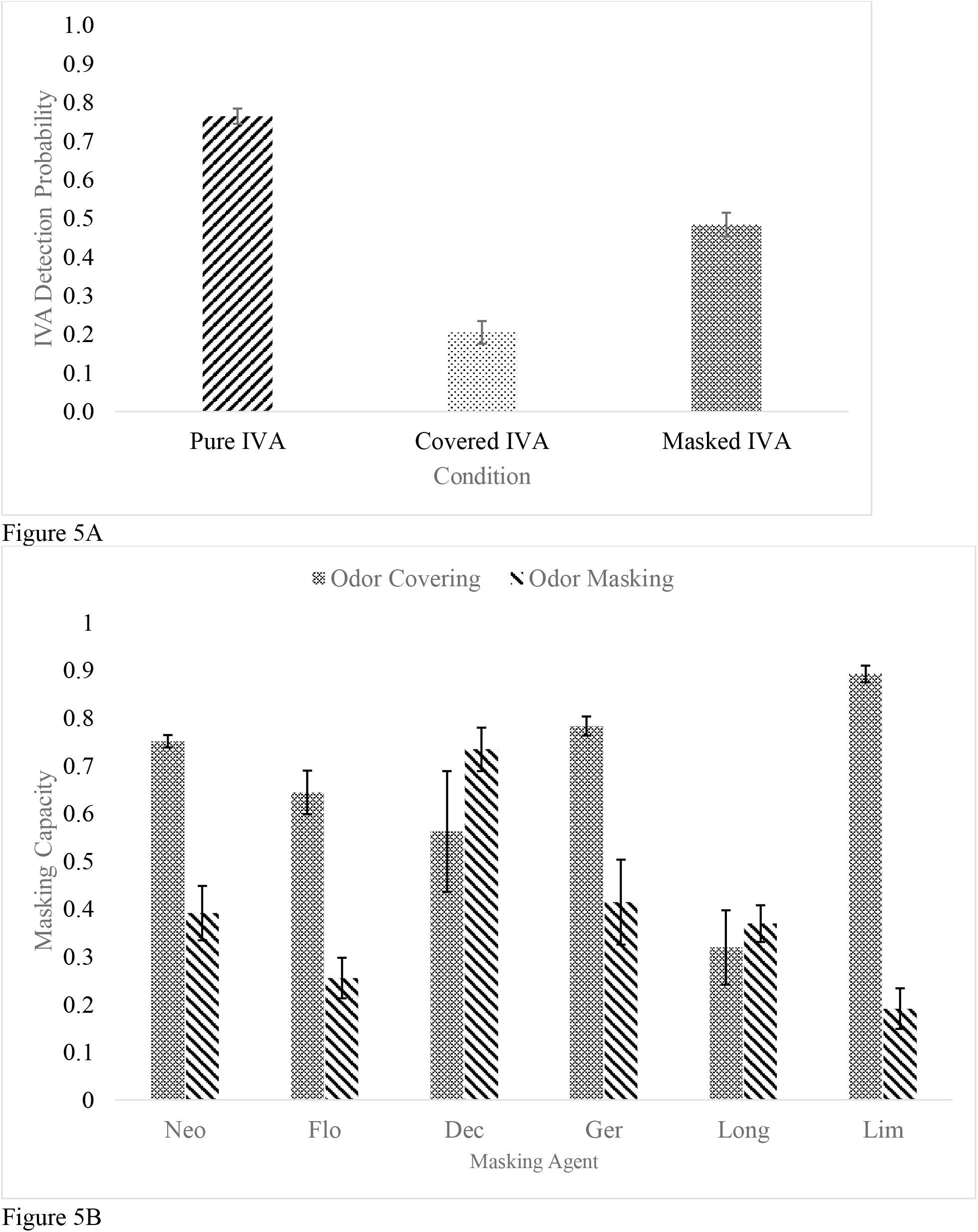
Figure 5A shows of bar graph of IVA detection probability when IVA is pure, covered (detectable amount of masking agents) and masked (undetectable amount of masking agents), which demonstrates that, in general, the odor covering effect contributes to more IVA detection probability drop than does odor masking effect. Figure 5B shows a Comparison of the IVA masking capacity for odor covering and odor masking of each masking agent across all concentrations used, which shows except for decanal and Iso-longifolanone, all other making agents possess stronger masking capacity when detectable.

### Short-term Habituation VS Long-term Habituation

Habituation has been defined as a decreased behavioral response to odors after repeated exposure to those odors (Pellegrino et al., 2017). In a recent review (Rankin et al., 2009), habituation was redefined as short-term habituation and long-term habituation based on the duration of habituation. We define short-term habituation as response decrement that could be restored in minutes (Philpott et al., 2008), while Long-term habituation as response decrement that “lasts hours, days or weeks” (P. Dalton, 1996). In our experiment, each visit includes 2 trials of the same odor stimulus. The IVA detection probabilities under all conditions between these 2 trials was plotted against each other (Figure 6B) and 2-way ANOVA was conducted to see if there was any significance between 2 trials. The IVA detection probability for the pure IVA possesses a subtle difference with P-adj = 0.076, while the other conditions showed no difference (P-adj > 0.05). These results indicate that the unpleasant odor, unmasked IVA, might have triggered a faster habituation onset, showing a potential relationship between hedonic level of odor and habituation onset time. The whisker plot of IVA detection probability for the pure IVA at each visit was shown in Figure 6A, and ANOVA was conducted to see if there is any significant difference. From our observation, there wasn’t any significant difference in IVA detection probability for the first 4 visits. Even the previous results showed there was likely a short-term habituation, the IVA sensitivity was always restored for the next visit, while at the 5th visit, the IVA detection dropped significantly and maintained at the same level at the 6th visit. IVA 5 and IVA 6 were at least 48 hrs apart from each other, meaning that the decrement in response was not restored to the previous level after the 5th session, suggesting long-term habituation. The same sample preparation procedure was used to prep the odorants during the experiments, suggesting the difference is not likely caused by the sample concentration difference or experimental error. When adjusting the IVA concentration to 10 PPM and 15 PPM respectively for Group 1 and Group 2, the IVA detection probability returned to 70% (Figure 3B), further suggesting that it was the habituation that caused the steep drop of IVA detection probability in the 5th session. It is still unclear when the habituation happened and why it happened that way.

**Figure 6.**
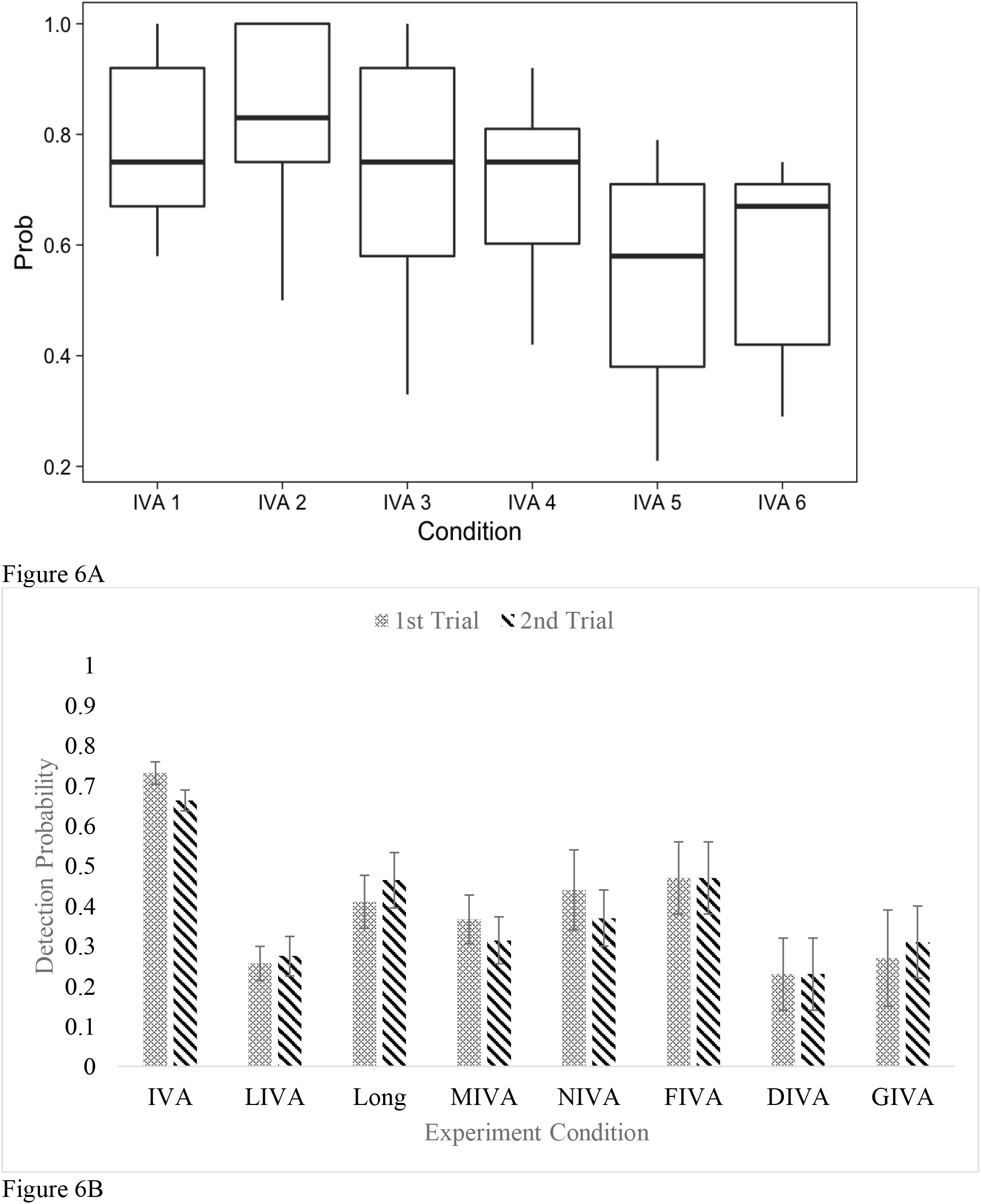
In Figure 6A, a boxplot of IVA 1-4 was plotted. The adjusted P-value for ANOVA when comparing IVA5 with IVA 1-4 is 0.003, 0.0003, 0.03, and 0.03 respectively, while the adjusted P-value for ANOVA between IVA 5 and IVA 6 is 0.78, demonstrating the IVA habituation started to happen at 5^th^ visit. In Figure 6B, IVA Detection probability for the 1^st^ trial and 2^nd^ trial that are 1 min apart from 1^st^ trial under each condition, which shows no significance short-term habituation happened between 2 experiment sessions.

### Masking effect of Masking Agents Mixture

With single masking agents, IVA could be masked to a level where it was unrecognizable (P < 0.5) (Figure 2B, Figure 2D), but only in rare cases did IVA became undetectable (P = 0). However, with 1.2PPM of scentless masking agent mixture, IVA detection probability dropped to 0.03 on average. Two theories could contribute to explaining this phenomenal masking effect by mixtures. First of all, masking could be achieved through competitive binding as was shown to play an inhibitory role during in-vitro ORs studies (Oka, Nakamura, et al., 2004; Oka, Omura, et al., 2004; Sanz et al., 2005) and in creating models that predicts odor mixture quality (Rospars et al., 2008; Singh et al., 2019). Theoretically, by using a peri-threshold level of masking agents’ mixture, masking agents will competitively bind to OR51E1, causing the activation to be minimized. Similar to previous studies (KAO et al., 2019), the antagonist against OR51E1 in our study, florhydral, demonstrated odor masking effect against IVA. However, the competitive binding theory on single OR might fail to explain *s-* limonene, Decanal, and Geraniol, which are odorants that does not bind to OR51E1 but still showed potent masking effect against IVA. Possible explanations for this phenomenon might be: 1) The In-vitro OR experimental procedure did not have the resolution to identify these PRMs as substrates, while they are antagonists or partial agonist to OR51E1; 2) OR51E1 not being the only major receptor for IVA, and these odorants are antagonistic to other unknown major receptors of IVA.

Another possible explanation comes from the odor “primacy code” theory, which hypothesizes the odor identification primarily relies on the activation pattern of a small set of earliest activated receptors (< 100ms) (Chong et al., 2020; Cleland et al., 2012; Wilson et al., 2017). In an extreme case, giving equal intensities of 30 odorants across olfactory space that potentially stimulate all receptors will render in an “Olfactory White” smell that does not contain characteristic of any its components (Weiss et al., 2012a). In our case, disrupting primacy code by using sub-threshold or peri-threshold amounts of odorants could likely make the malodor, *iso*-valeric acid in this case, unrecognizable.

## Conclusion

In our experiment, we were able to demonstrate that peri-threshold amounts of single PRMs were able to mask the smell of isovaleric acid, and the effectiveness of the masking follows a dose dependent pattern with the amount of masking agents within a critical range. We further demonstrated that, by using a mixture of peri-threshold amount of masking agents, a complete masking of IVA could be achieved while the masking agents remain undetectable. Future research will be focusing on two aspects: 1) studying if the odor masking is unilateral (only masking agents can mask IVA) or mutual (IVA and masking agents masking each other); 2) finding a universal masking agent mixture that could potentially mask the majority of malodors.

## Ethics Statement

This protocol (Protocol ID#: 912009302) has been assessed and approved by the Institutional Review Board for Human Participant Research at Cornell University. All participants read and signed an IRB approved consent form prior to participation.

## Acknowledgement

I would like to thank Matt Wagner, Marisa Casola and Jennifer Nienaber from Procter & Gamble Co. for supporting the funding of this research by Procter & Gamble Co..

