## Supplementary material for "Masking effects on *Iso*-valeric Acid Recognition by Sub-threshold Odor Mixture": Figures

Figure 2A.

| | IVA Average | NIVA | FIVA | DIVA | GIVA | Long | MIVA1 | LIVA 1 | LIVA 2 |
|---|---|---|---|---|---|---|---|---|---|
| Mean | 0.76 | 0.4 | 0.47 | 0.23 | 0.29 | 0.45 | 0.25 | 0.11 | 0.09 |
| Std | 0.17 | 0.25 | 0.26 | 0.26 | 0.32 | 0.32 | 0.27 | 0.2 | 0.22 |
| SE | 0.02 | 0.06 | 0.06 | 0.06 | 0.07 | 0.08 | 0.06 | 0.05 | 0.05 |
| SE UP | 0.01 | 0.03 | 0.03 | 0.03 | 0.035 | 0.04 | 0.03 | 0.025 | 0.025 |
| SE Down | 0.01 | 0.03 | 0.03 | 0.03 | 0.035 | 0.04 | 0.03 | 0.025 | 0.025 |

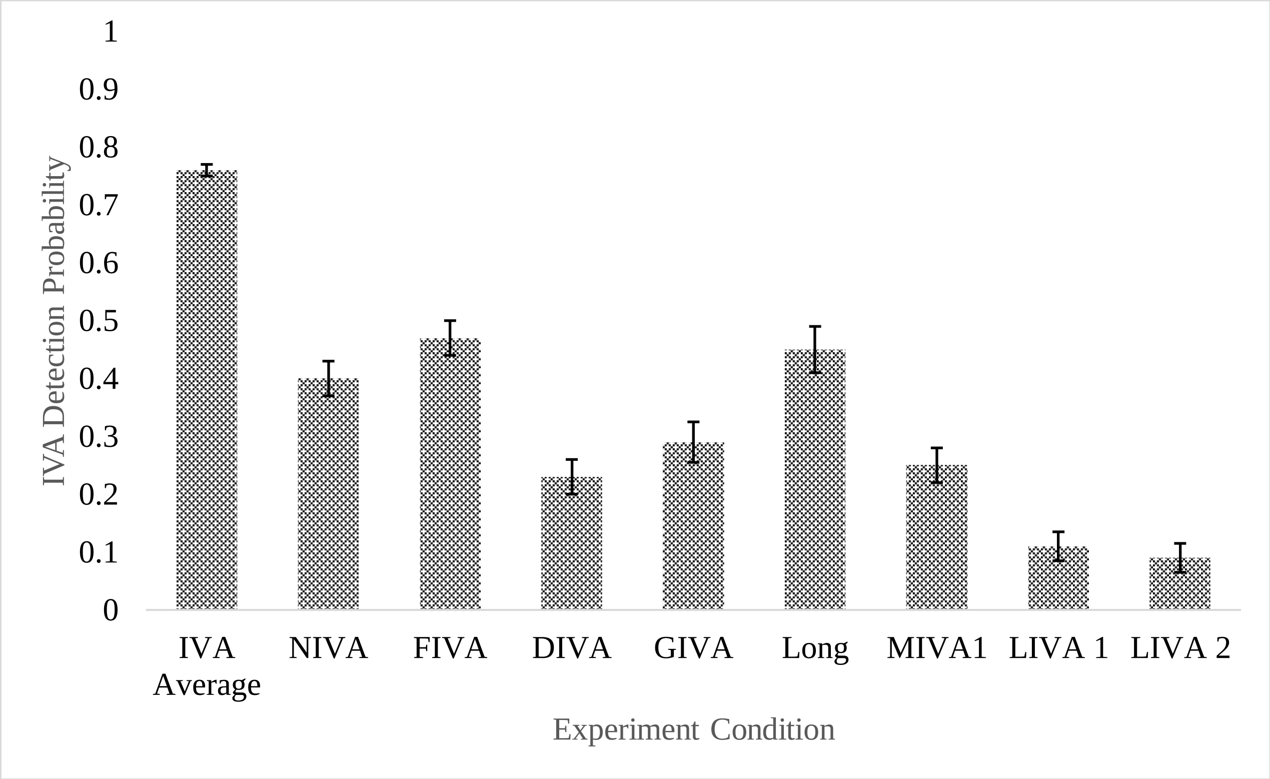

Figure 2D.

| | IVA Average | MIVA 2 | LIVA 3 | Long 2 | LIVA 4 |
|---|---|---|---|---|---|
| Average | 0.56 | 0.43 | 0.5 | 0.4 | 0.38 |
| Std | 0.2 | 0.2 | 0.2 | 0.23 | 0.23 |
| SE | 0.03 | 0.05 | 0.06 | 0.05 | 0.05 |
| SE up | 0.015 | 0.025 | 0.03 | 0.025 | 0.025 |

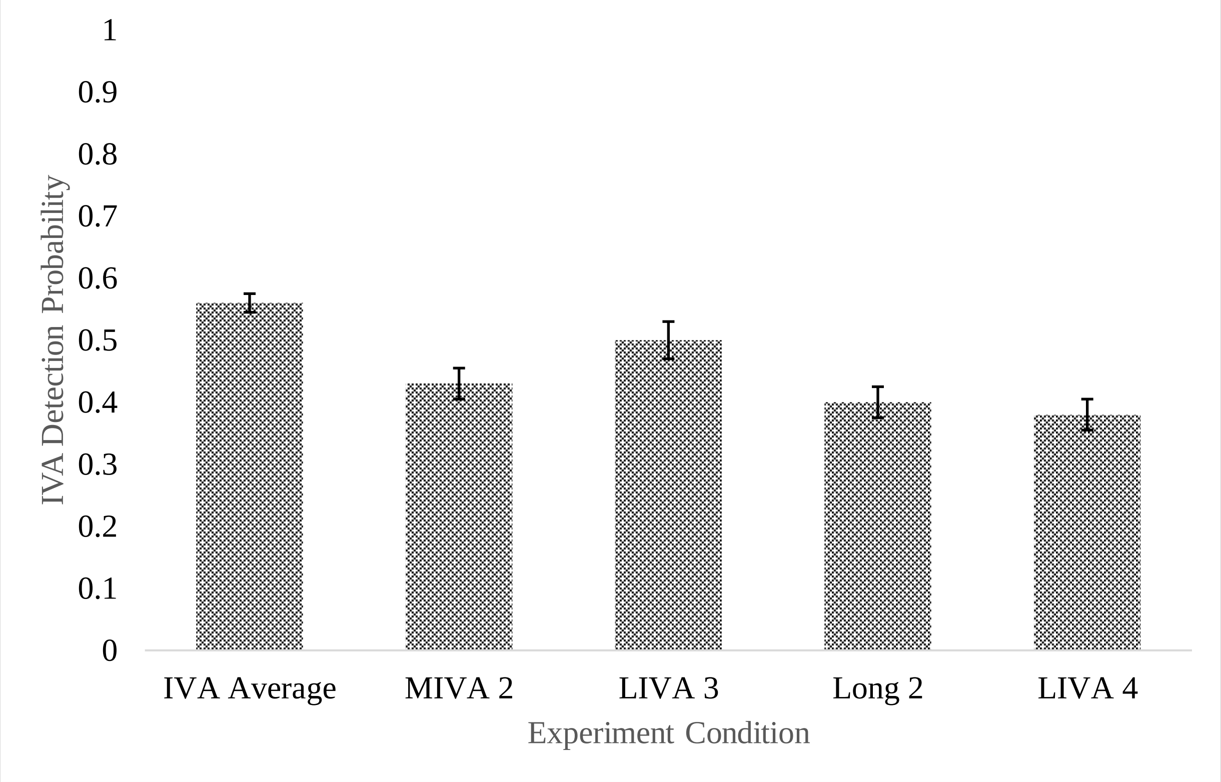

Figure 3B.

| | Blank | IVA | Sum 1.2PPM |
|---|---|---|---|
| Subject 1 | 0 | 0.92 | 0 |
| | 0 | 1 | 0.08 |
| Subject 2 | 0 | 0.92 | 0 |
| | 0 | 0.67 | 0 |
| Subject 3 | 0.08 | 0.75 | 0 |
| | 0 | 0.67 | 0 |
| Subject 4 | 0.08 | 1 | 0.08 |
| | 0.08 | 0.92 | 0 |
| Subject 5 | 0 | 0.92 | 0.25 |
| | 0 | 0.58 | 0.17 |
| Subject 6 | 0 | 0.83 | 0 |
| | 0 | 0.5 | 0 |
| Subject 7 | 0 | 0.67 | 0 |
| | 0 | 0.25 | 0 |
| Subject 8 | 0.08 | 0.58 | 0 |
| | 0 | 0.67 | 0 |
| Subject 9 | 0 | 0.75 | 0 |
| | 0 | 0.67 | 0 |

| | Blank | IVA | IVA + Masking Mixture |
|---|---|---|---|
| Average | 0.02 | 0.74 | 0.03 |
| STD | 0.034 | 0.195 | 0.071 |
| SE | 0.008 | 0.046 | 0.017 |
| SE up | 0.00403 | 0.02298 | 0.00836 |
| SE down | 0.00403 | 0.02298 | 0.00836 |

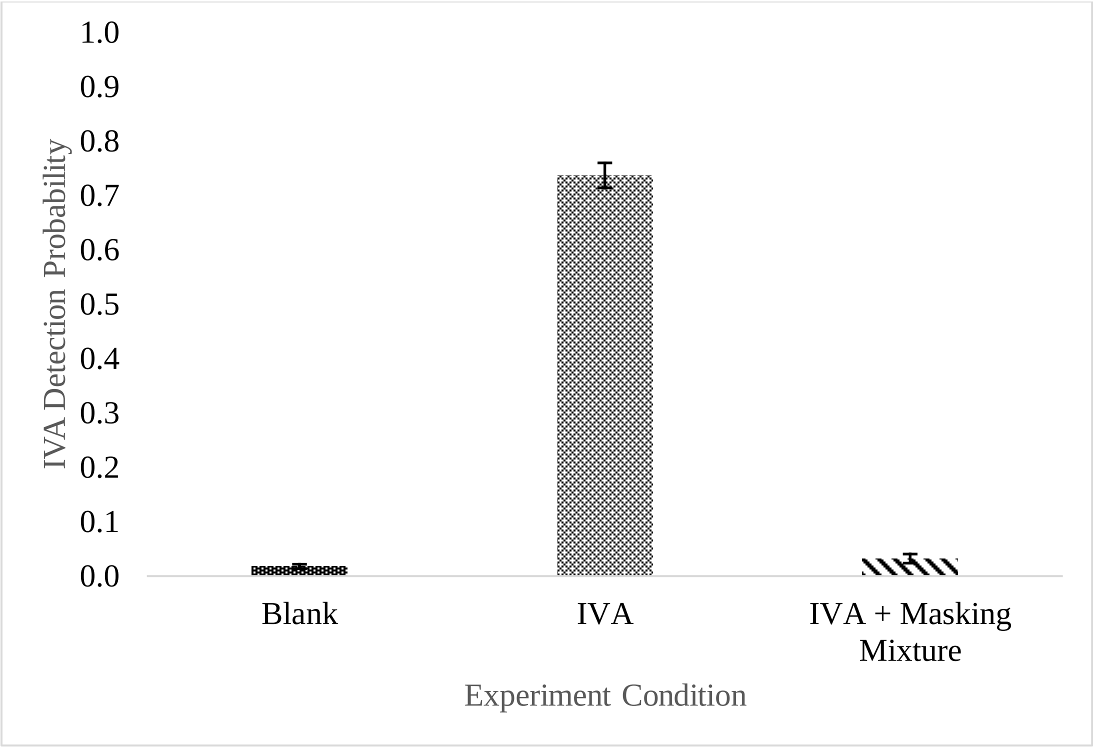
1.0
0.9
0.8
0.7
0.6
0.5
0.4
0.3
0.2
0.1
0.0
IVA Detection Probability
Blank
IVA
IVA + Masking Mixture
Experiment Condition

Figure 4A.

| | Neo | Flo | Dec | Ger | Met | Long | Lim |
|---|---|---|---|---|---|---|---|
| Masking Capacity | 0.494 | 0.415 | 0.719 | 0.605 | 0.143 | 0.274 | 0.292 |
| SE/2 | 0.087 | 0.081 | 0.1 | 0.131 | 0.139 | 0.139 | 0.13 |
| Dose | 0.467 | 0.107 | 0.334 | 0.322 | 0.5 | 0.3 | 0.05 |
| SE/2 | 0.044 | 0.028 | 0.03 | 0.037 | 0 | 0 | 0 |

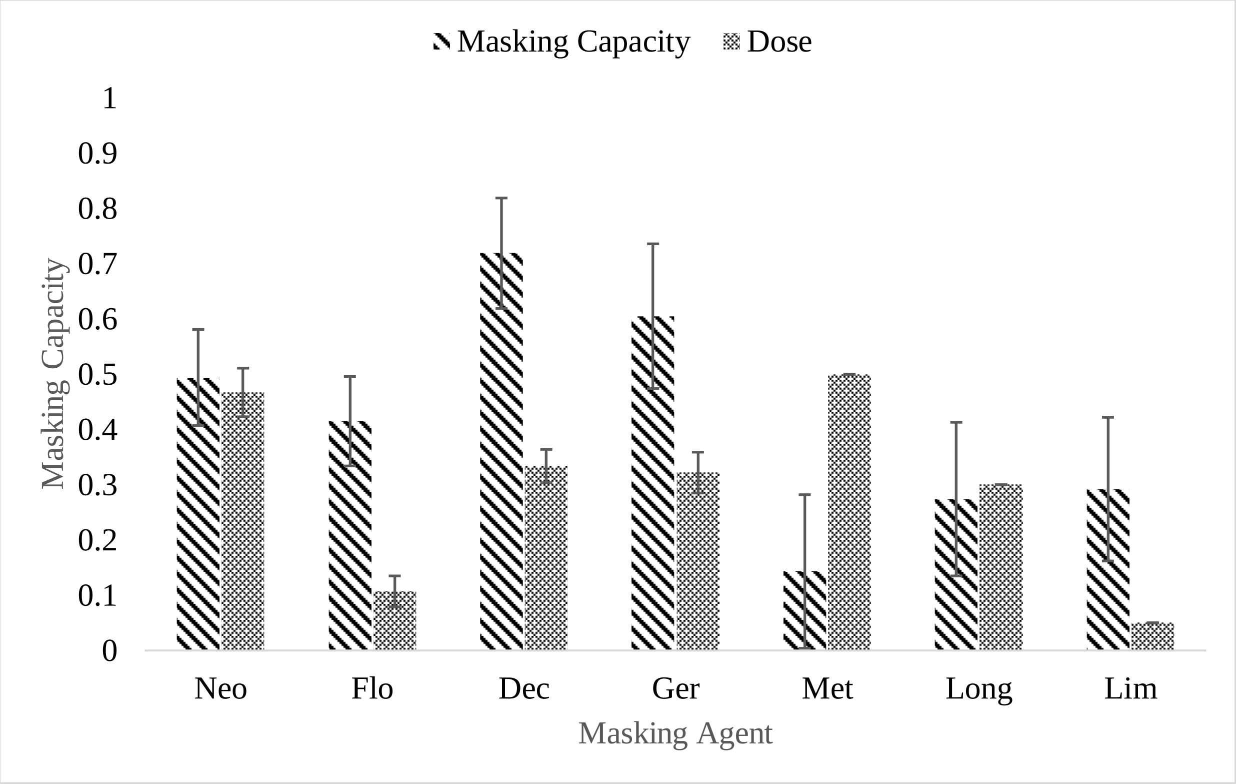

Figure 4B.

| | Dose | Masking Capacity |
|---|---|---|
| Neo | 0.467 | 0.494 |
| Flo | 0.107 | 0.415 |
| Dec | 0.334 | 0.719 |
| Ger | 0.322 | 0.605 |
| Long | 0.3 | 0.274 |
| Lim | 0.05 | 0.292 |

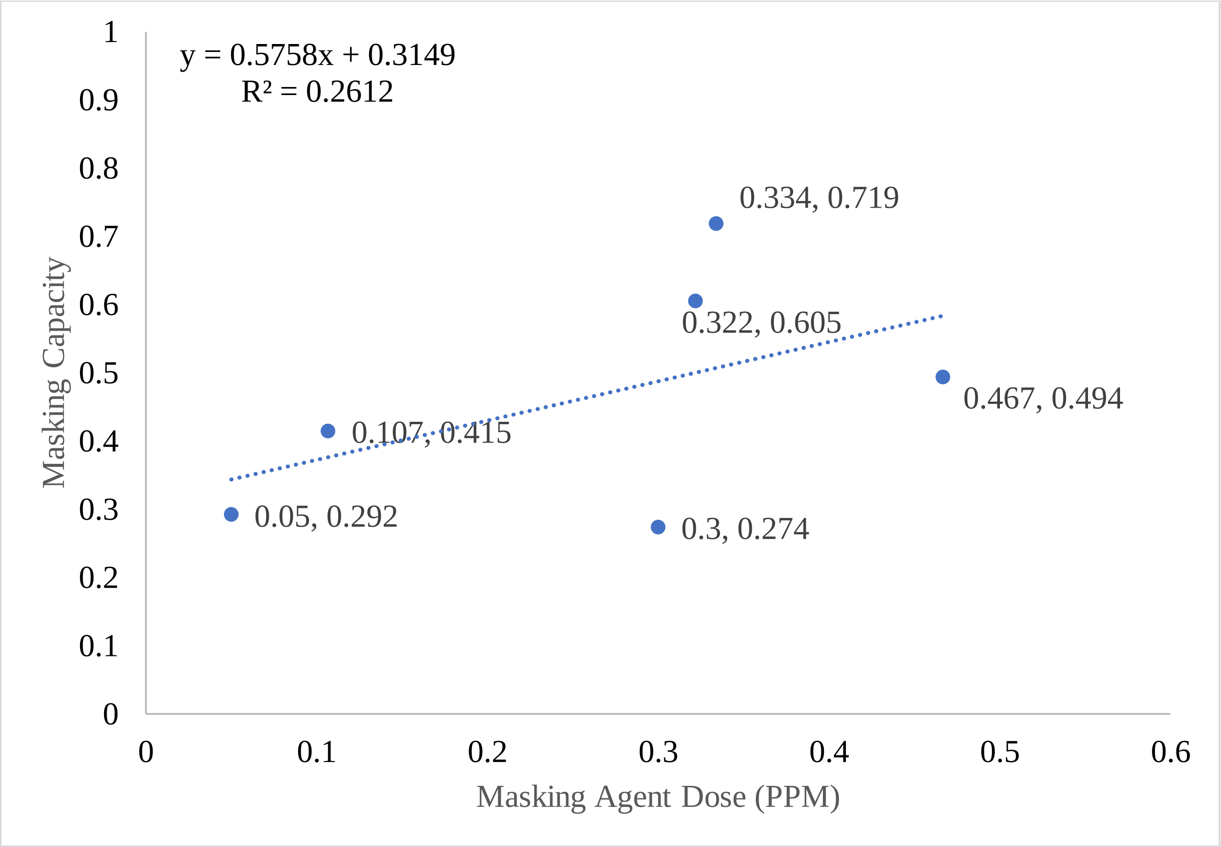

Figure 4C.

| | | | | |
|---|---|---|---|---|
| Concentration | 2.65 | 1 | 0.05 | 0.02 |
| Masking Ratio | 0.87 | 0.87 | 0.29 | 0.13 |

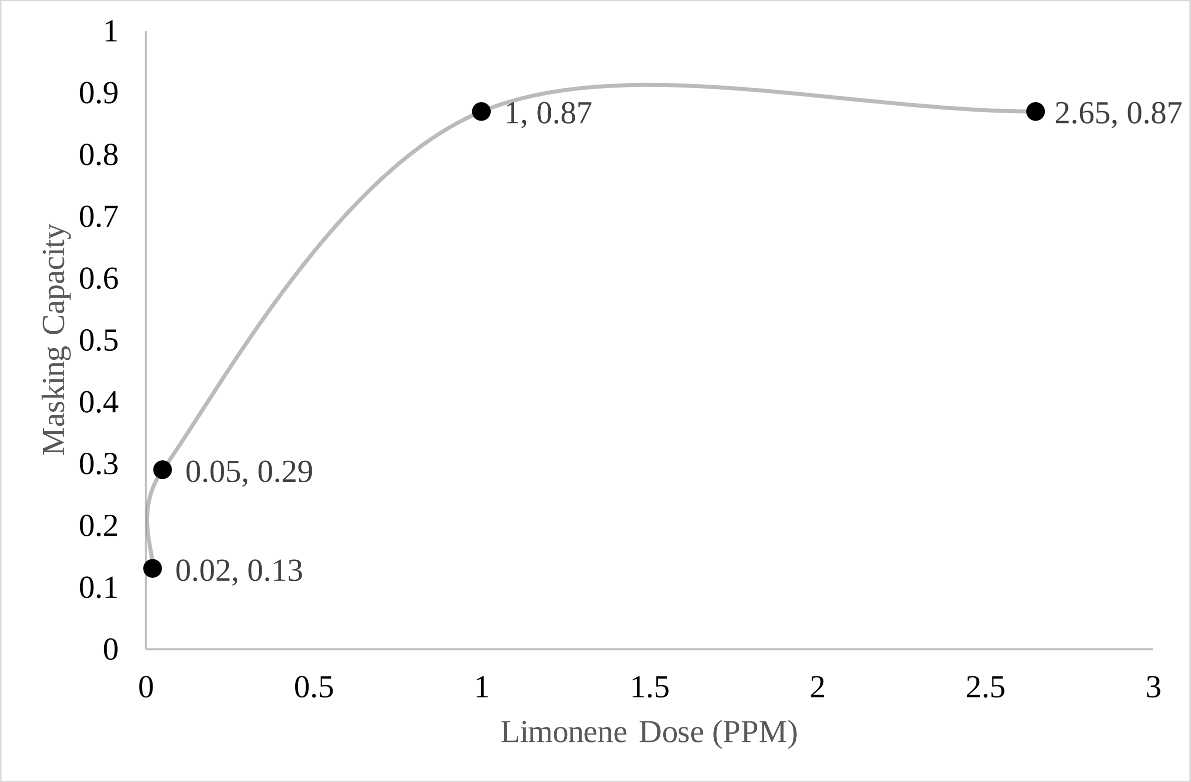

Figure 5A.

| | Pure IVA | Covered IVA | Masked IVA |
|---|---|---|---|
| Average | 0.764 | 0.205 | 0.484 |
| SE | 0.020 | 0.029 | 0.031 |

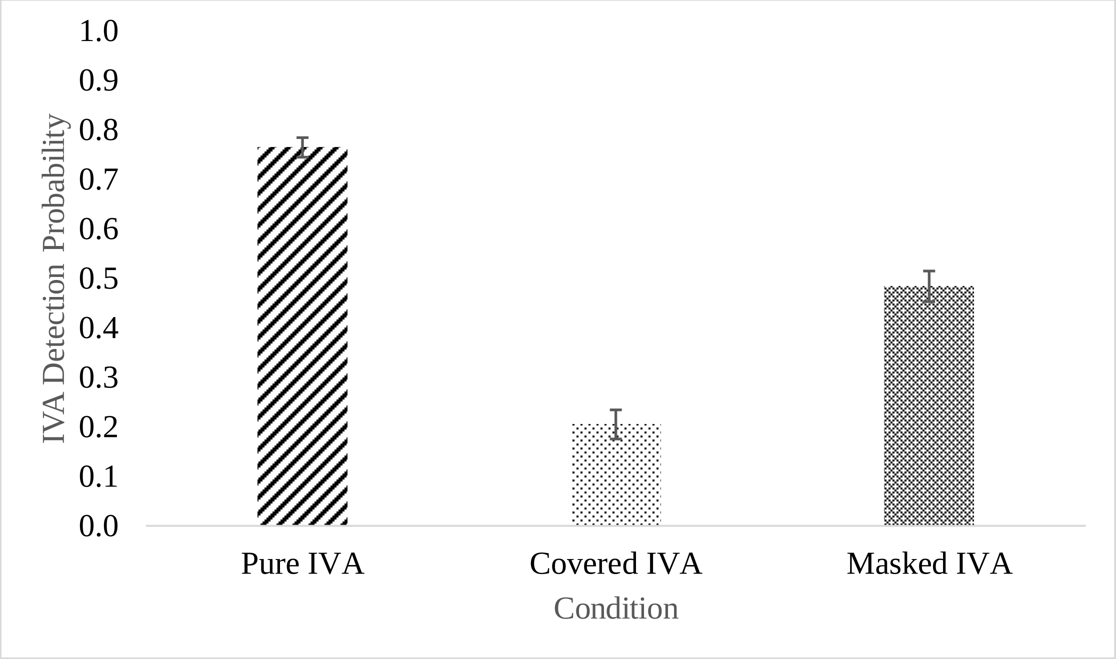

Figure 5B.

| | Neo | Flo | Dec | Ger | Long | Lim |
|---|---|---|---|---|---|---|
| Odor Covering | 0.751 | 0.644 | 0.562 | 0.783 | 0.319 | 0.892 |
| SE | 0.026 | 0.091 | 0.253 | 0.04 | 0.156 | 0.035 |
| SE Up | 0.013 | 0.0455 | 0.1265 | 0.02 | 0.078 | 0.0175 |
| SE Down | 0.013 | 0.0455 | 0.1265 | 0.02 | 0.078 | 0.0175 |
| Odor Masking | 0.391 | 0.255 | 0.734 | 0.414 | 0.369 | 0.191 |
| SE | 0.114 | 0.085 | 0.091 | 0.178 | 0.077 | 0.085 |
| SE Up | 0.057 | 0.0425 | 0.0455 | 0.089 | 0.0385 | 0.0425 |
| SE Down | 0.057 | 0.0425 | 0.0455 | 0.089 | 0.0385 | 0.0425 |

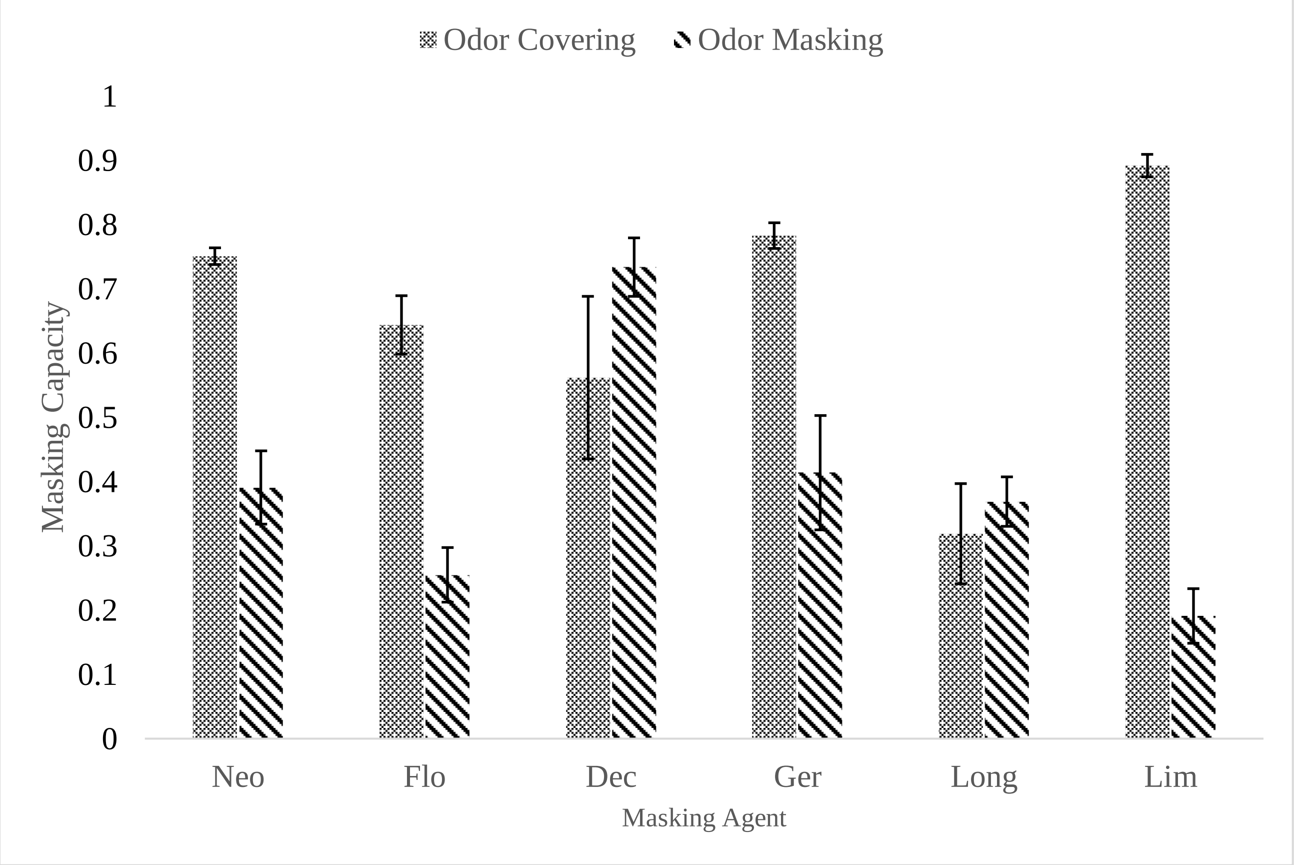
