## Supplemental tables for "Masking effects on *Iso*-valeric Acid Recognition by Sub-threshold Odor Mixture"

*Figure 1: Figure 1A shows 3 bottles containing ascending concentrations of an odorant were labeled from 1 to 3. In one trial, each bottle was puffed 4 times randomly. For each experiment session, total of 3 trials were conducted, so that each bottle was puffed 4 times at each position on the triad shown in Figure 1B*

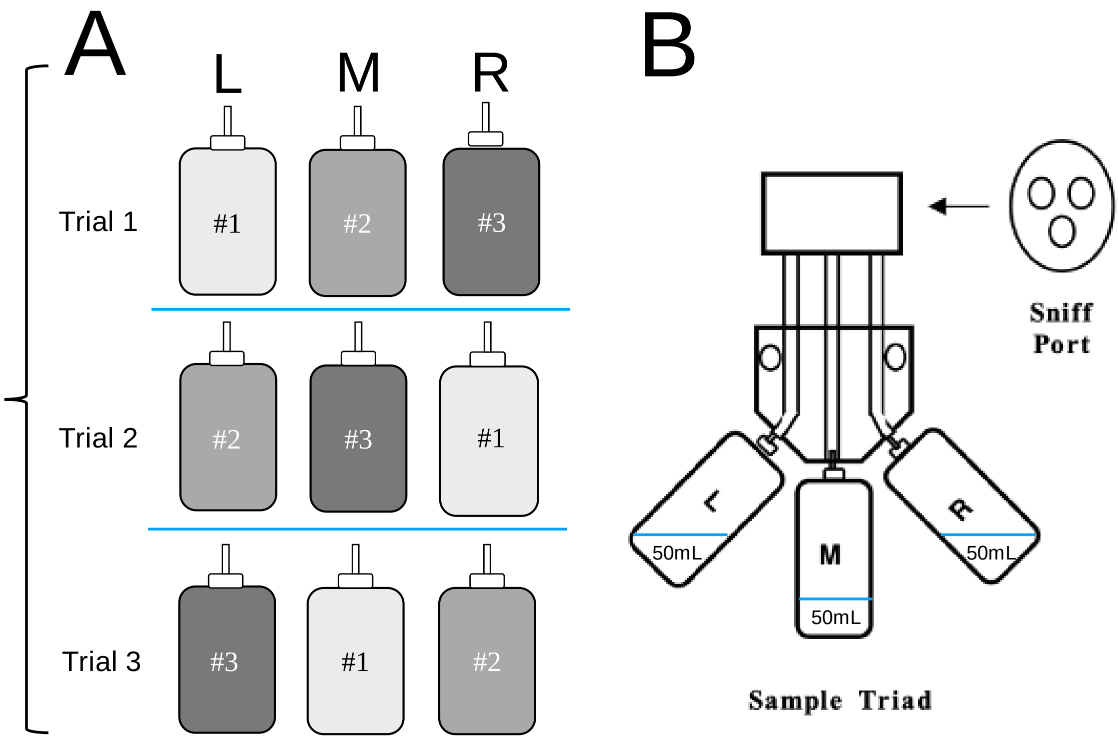


*Table 1. Concentrations and log of concentrations of each PRMs used for threshold determination.*

| Odorants | Concentrations (PPM) | | | Log of Concentrations | | |
|---|---|---|---|---|---|---|
| **IVA** | 0.5 | 5 | 20 | -0.3 | 0.7 | 1.3 |
| **Neohivernal** | 0.1 | 1 | 5 | -1 | 0 | 0.7 |
| **Decanal** | 0.01 | 0.5 | 3 | -2 | -0.3 | 0.5 |
| **Florhydral** | 0.01 | 0.05 | 0.1 | -2 | -1.3 | -1 |
| **Geraniol** | 0.01 | 0.3 | 2.5 | -2 | -0.5 | 0.4 |
| **Methyl *Iso*-eugenol** | 5 | 30 | 80 | 0.7 | 1.5 | 1.9 |
| ***Iso*-longifolanone** | 1 | 10 | 30 | 0 | 1 | 1.5 |
| ***Iso*-longifolanone (2)** | 0.1 | 1 | 25 | -1 | 0 | 1.4 |
| ***S*-Limonene** | 1 | 10 | 30 | 0 | 1 | 1.5 |

*Figure 2: Figure 2 shows the psychometric curve of IVA detection threshold generated using 3 concentrations of IVA for subject 4. The recognition threshold was determined by the concentration at which the detection probability reaches 0.5.*

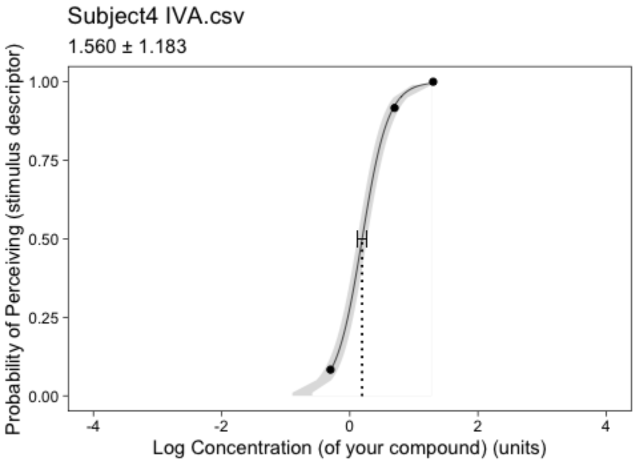


*Table 2. Measured recognition thresholds of 8 odorants for subjects. ND (Not Detectable) means this subject could not smell the odorant at the tested concentrations.*

| Subject / Threshold (PPM) | IVA | Neo | Flo | Dec | Long | Ger | Met | Lim |
|---|---|---|---|---|---|---|---|---|
| **Subject 1** | 0.40 | 1.03 | 0.30 | 0.54 | 6.51 | 0.37 | 72.86 | 3.24 |
| **Subject 2** | 0.49 | 1.19 | 0.02 | 0.44 | 10.69 | 0.39 | 5.25 | 8.9 |
| **Subject 3** | 1.00 | 0.21 | 0.02 | 0.48 | 0.27 | 0.34 | 29.17 | 3.24 |
| **Subject 4** | 1.60 | 2.00 | 0.07 | 0.70 | 3.77 | 0.34 | 18.53 | 1.15 |
| **Subject 5** | 2.60 | 1.26 | 0.07 | 1.21 | 23.29 | 1.40 | 80.00 | 5.58 |
| **Subject 6** | 4.90 | 0.63 | 0.67 | 1.00 | 9.46 | 0.57 | 31.83 | 9.5 |
| **Subject 7** | 5.00 | 1.01 | 0.02 | 0.48 | 4.42 | ND | 15.64 | 4.1 |
| **Subject 8** | 5.30 | 0.28 | 0.02 | 0.02 | 2.44 | 0.59 | 27.73 | 3.09 |
| **Subject 9** | 7.70 | 1.01 | 0.07 | 1.21 | 10.28 | 0.16 | 32.18 | 9.27 |
| **Average** | 3.05 | 1.06 | 0.14 | 0.72 | 7.19 | 0.56 | 35.36 | 5.78 |
