## Supplementary material for "Masking effects on *Iso*-valeric Acid Recognition by Sub-threshold Odor Mixture": R code

**Section 1: Psychometric Curve Generation for Recognition Threshold**

```r
# File uploading
filename <- File

# Data Cleaning
label <- tail(strsplit(filename,split="/")[[1]],1)
data <- read.csv(filename)
data_n<- data %>% group_by(LogConR) %>%
  count(LogConR)
N <- data_n[,2]
data_c<- data %>% group_by(LogConR) %>%
  summarise(Success = sum(Resp))%>%
  cbind(N)%>%
  mutate(Failure =n-Success, Prob=Success/n)

#Modeling
m1<-glm(Resp~ LogConR, data=data, family="binomial")

m2<-glm(cbind(Success, Failure)~ LogConR, data=data_c,
family="quasibinomial")
summary(m2)

calibrate <-expand.grid(Prob=seq(0.05,0.95,0.01))
calibrate$LogConR<-NA
calibrate$se<-NA
for (i in 1:nrow(calibrate)){
  #list out attribute, dose-response curve based
  calibrate$se[i]<-attributes(dose.p(m2, cf=1:2, p=calibrate$Prob[i]))$SE[,1]
  calibrate$LogConR[i]<-dose.p(m2, cf=1:2, p=calibrate$Prob[i])[1]
}

#making variables for data visualization
data_d <- calibrate
data_d$min <- NA
data_d$max <- NA
for (i in 1:nrow(data_d)){
  data_d$min[i] <- (data_d$LogConR[i] - data_d$se[i])
  data_d$max[i] <- (data_d$LogConR[i] + data_d$se[i])
}

#get two more model entries closer to zero and one

dose_a <- dose.p(m2, cf=1:2, p=0.01)
a1 <- dose_a[1]
dose_aSE <- data.frame(attributes(dose_a))
a2 <- dose_aSE[2]

dose_b <-dose.p(m2, cf=1:2, p=0.99)
```

```r
b1 <- dose_b[1]
dose_bSE <- data.frame(attributes(dose_b))
b2 <- dose_bSE[2]

data_e <- rbind(data_d, list(0.01, a1, a2, (a1 - a2), (a1 + a2)))
data_e <- rbind(data_e, list(0.99, b1, b2, (b1 - b2), (b1 + b2)))

data_e$LogConR <- as.numeric(data_e$LogConR)
data_e$min <- as.numeric(data_e$min)
data_e$max <- as.numeric(data_e$max)
data_e$se <- as.numeric(data_e$se)

f <- data_e[46,2] #log of the threshold
g <- 10^f #take it back to actual concentrations
g <- roundString(g, 3) #round to 3 decimal places
h <- roundString(10^(1.96*(data_e$se[46])), 3) #make confidence interval
#g is Prob = 0.5 point
#h is 95% confidence interval at P = 0.5
v <- "\u00B1" #unicode symbol for plus/minus sign
g2 <- paste(g, substr(v, 1, 1), h) #threshold plus/minus confidence interval

#adding one more modeling row
data_f <- data_e
w <- data_f[93, 1] #prob
e <- data_f[93, 2] #LogConR
r <- data_f[93, 3] #se
t <- data_f[93, 5] #max
data_f <- rbind(data_f, list(w, e, r, t, t))

# Plotting
ggplot()+
   geom_area(data_f, mapping = aes(x = (2*(min - LogConR) + LogConR), y =
Prob), fill = "grey", alpha = 0.5)+
  geom_area(data_e, mapping = aes(x = (2*(max - LogConR) + LogConR), y =
Prob), fill = "white")+
  geom_point(data = data_c, aes(x=LogConR, y=Prob))+
  geom_smooth(data = data_c, aes(x=LogConR, y=Prob), method = "glm",
method.args = list(family="quasibinomial"), color = "black", size = 0.25, se
= F) +
  geom_errorbarh(data=data_e, aes(y=Prob[46], xmax=LogConR[46]+2*se[46],
xmin=LogConR[46]-2*se[46]), height=0.05, size= 0.125) +
  geom_segment(data = data_c, aes(x = f, y = 0, xend = f, yend = 0.5),
linetype = "dotted")+

  # x axis restriction. coord_cartesian restrict the range that shown
  coord_cartesian(xlim = c(-3,4)) +
  # graph labels
  ggtitle(label, subtitle = g2)+
  xlab("Log Concentration (of your compound) (units)") +
```

```
  ylab("Probability of Perceiving (stimulus descriptor)")+

  #cosmetic changes below here
  theme_linedraw()+
  theme(panel.background = element_rect(fill = "white"),
        panel.grid.major = element_blank(),
        panel.grid.minor = element_blank())
```

**Section 2: Anova Analysis for Habituation**

```
overall = read.csv(File)

#Data Filtering
Compared_data <- overall %>%
  filter(Condition == "IVA 1" | Condition == "IVA 2"| Condition == "IVA 3"|
Condition == "IVA 4"|Condition == "IVA 5"|Condition == "IVA 6")

# Modeling
Anova.model <- aov(data = Compared_data, Prob ~ Condition)
summary(Anova.model)

# Plot
ggplot()+
  geom_boxplot(data = Compared_data, mapping = aes(x = Condition, y = Prob))+
  theme_linedraw()+
  theme(panel.background = element_rect(fill = "white"),
        panel.grid.major = element_blank(),
        panel.grid.minor = element_blank())
```
